# Inhibition of HtrA2 alleviated colitis by preventing necroptosis of intestinal epithelial cells

**DOI:** 10.1101/395897

**Authors:** Chong Zhang, Andong He, Shuai Liu, Qiaoling He, Yiqin Luo, Yujiao Chen, Zhilan He, Ailin Tao, Jie Yan

## Abstract

Necroptosis of intestinal epithelial cells has been indicated to play an important role in the pathogenesis of inflammatory bowel disease (IBD). The identification of dysregulated proteins that can regulate necroptosis in dextran sulfate sodium (DSS)-induced colitis is the key to the rational design of therapeutic strategies for colitis. Through Tandem Mass Tag (TMT)-based quantitative proteomics, HtrA2 was found to be downregulated in the colon of DSS-treated mice. UCF-101, a specific serine protease inhibitor of HtrA2, significantly alleviated DSS-induced colitis as indicated by prevention of body weight loss and decreased mortality. UCF-101 decreased DSS-induced colonic inflammation, prevented intestinal barrier function loss and inhibited necroptosis of intestinal epithelial cells. *In vitro*, UCF-101 or silencing of HtrA2 decreased necroptosis of HT-29 and L929 cells. UCF-101 decreased phosphorylation of RIPK1 and subsequent phosphorylation of RIPK3 and MLKL during necroptosis. HtrA2 directly interacted with RIPK1 and promoted its degradation during a specific time phase of necroptosis. Our findings highlight the importance of HtrA2 in regulating colitis by modulation of necroptosis and suggest HtrA2 as an attractive target for anti-colitis treatment.

## Introduction

Inflammatory bowel diseases (IBD), namely Crohn’s disease and ulcerative colitis, affect about 3.7 million people in the United States and Europe (Ananthakrishnan, 2015). However, its etiology remains elusive, as complex interactions between genetic susceptibility, microbial dysbiosis, and environmental factors are involved in its pathogenesis (Maloy et al, 2011). Intestinal epithelium provides a physical barrier that modulates microbial colonization and prevents their penetration of the epithelium (Nowarski et al, 2015). Epithelial cell death is a hallmark of intestinal inflammation and leads to intestinal barrier disruption, which contributes to the pathogenesis of IBD (Luissint et al, 2016). Necroptosis, a newly recognized programmed cell death, of intestinal epithelial cells led to disruption of the intestinal barrier and resulted in spontaneous colitis or terminal ileitis in mice (Gunther et al, 2011; Welz et al, 2011). Intervening in necroptosis has been indicated as a promising therapeutic strategy for IBD. However, the role of deregulated genes that contribute to necroptosis in IBD remains largely unexplored.

Necroptosis is typically considered a highly pro-inflammatory mode of cell death, due to release of intracellular “damage-associated molecular patterns” that promote inflammation (Kearney et al, 2017). Necroptosis is a caspase-independent cell death and can be initiated by death receptors including TNFR1, TLR3, TLR4 and IFNRs (Kearney et al, 2017; Weinlich et al, 2017). Signal transduction during necroptosis has been well studied in the context of TNF-α. Upon TNF-α stimulation, RIPK1, FADD and CYLD are recruited to TNFR1 to form a protein complex. Subsequent deubiquitylation and phosphorylation events lead to RIPK1 phosphorylation and activation (de Almagro et al, 2017; Moquin et al, 2013). When caspase-8 activity is absent or inhibited, the phosphorylated RIPK1 regulates the formation of a necrosome which consists of RIPK1, RIPK3 and MLKL (Grootjans et al, 2017). Via RIP homotypic interaction motif-domain (RHIM) interactions, RIPK1 promotes oligomerization and subsequently autophosphorylation of RIPK3 (Orozco et al, 2014; Wu et al, 2014). MLKL is then recruited to the RIPK1/RIPK3 complex and phosphorylated by p-RIPK3. Phosphorylated MLKL forms oligomers and translocates to the intracellular plasma membrane where it binds to phosphatidylinositol lipids and cardiolipin, leading to the formation of pores and finally disrupting cellular membrane integrity (Wang et al, 2014a). As an essential factor for necroptosis in the context of TNF-α, RIPK1 is reported to regulate necroptosis positively in a kinase dependent manner, and negatively in a kinase independent manner whereby the scaffolding function of the RHIM domain prevents ZBP1 from activating RIPK3 and thus represses necroptosis (Dannappel et al, 2014; Newton et al, 2016b; Orozco et al, 2014).

HtrA2 is a serine protease located in mitochondria and involved in apoptosis regulation (Suzuki et al, 2001). Upon apoptotic stimuli, HtrA2 translocates from mitochondria to cytosol, where it binds to and cleaves IAPs, thus releasing caspases from their natural inhibitors (Yang et al, 2003). Independently, HtrA2 can cleave anti-apoptotic proteins ped/pea 15 and Hax-1 and thereby promote apoptosis (Liu et al, 2017; Trencia et al, 2004). In addition to apoptosis, HtrA2 is also reported to have a role in necroptosis. In IL-13 deprivation induced cell death, HtrA2 was able to cleave RIPK1 and enhanced cell death in a caspase independent manner (Vande Walle et al, 2010). According to the references, HtrA2 promotes necroptosis in a serine protease dependent manner (Blink et al, 2004; Sosna et al, 2013). UCF-101, a serine protease inhibitor of HtrA2, could inhibit TNF-α plus Z-VAD induced necroptosis in neutrophils, L929Ts and Jurkat I42 cells, as well as TNF-α plus Z-VAD and cycloheximide induced necroptosis in HT-29 cells (Blink et al, 2004; Sosna et al, 2013). Nonetheless, the exact mechanism for HtrA2 in necroptosis regulation and IBD pathogenesis still remains unknown and needs further investigation.

Based on high throughput proteomic analysis, we found significant downregulation of HtrA2 in colons of DSS treated mice. Moreover, the pathological symptoms and animal motility were ameliorated by UCF-101 treatment. Mechanism investigation suggested that HtrA2 could interact with RIPK1, promote its degradation and finally enhance necroptosis. Our study raises the possibility of HtrA2 as a potential therapeutic target for anti-colitis treatment.

## Results

### HtrA2 is downregulated in colons of DSS-treated mice

To identify important proteins that are dysregulated in the inflamed colon that could be used as novel potential therapeutic targets, we utilized a DSS-induced colitis mouse model for quantitative proteomics. Control mice were given distilled water for 10 days while DSS-treated mice were given 3% DSS for 7 days that was replaced with distilled water for the following 3 days. Based on daily monitoring, the DSS-induced mice slowly began to lose weight and have bloody stools. On day 7, the symptoms were most severe and some of the mice died. The surviving mice recovered gradually and eventually returned to normal. Thus, we harvested the colon tissues on day 7 and day 10 to detect protein levels by TMT quantitative proteomics. Compared with the distilled water-treated control mice, HtrA2 protein levels were significantly decreased in the colons of DSS-treated mice (Fig EV1 and Fig 1A). In addition, the downregulation of HtrA2 in the colons of DSS-treated mice was confirmed by immunoblotting and immunohistochemical (IHC) staining (Fig 1B-1C). These data show that the protein level of HtrA2 was significantly reduced as colitis progressed. Further study is needed to determine whether the downregulation of HtrA2 is the cause of colitis development or whether the negative feedback protective mechanism is initiated by the colon tissue.

**Figure 1.**
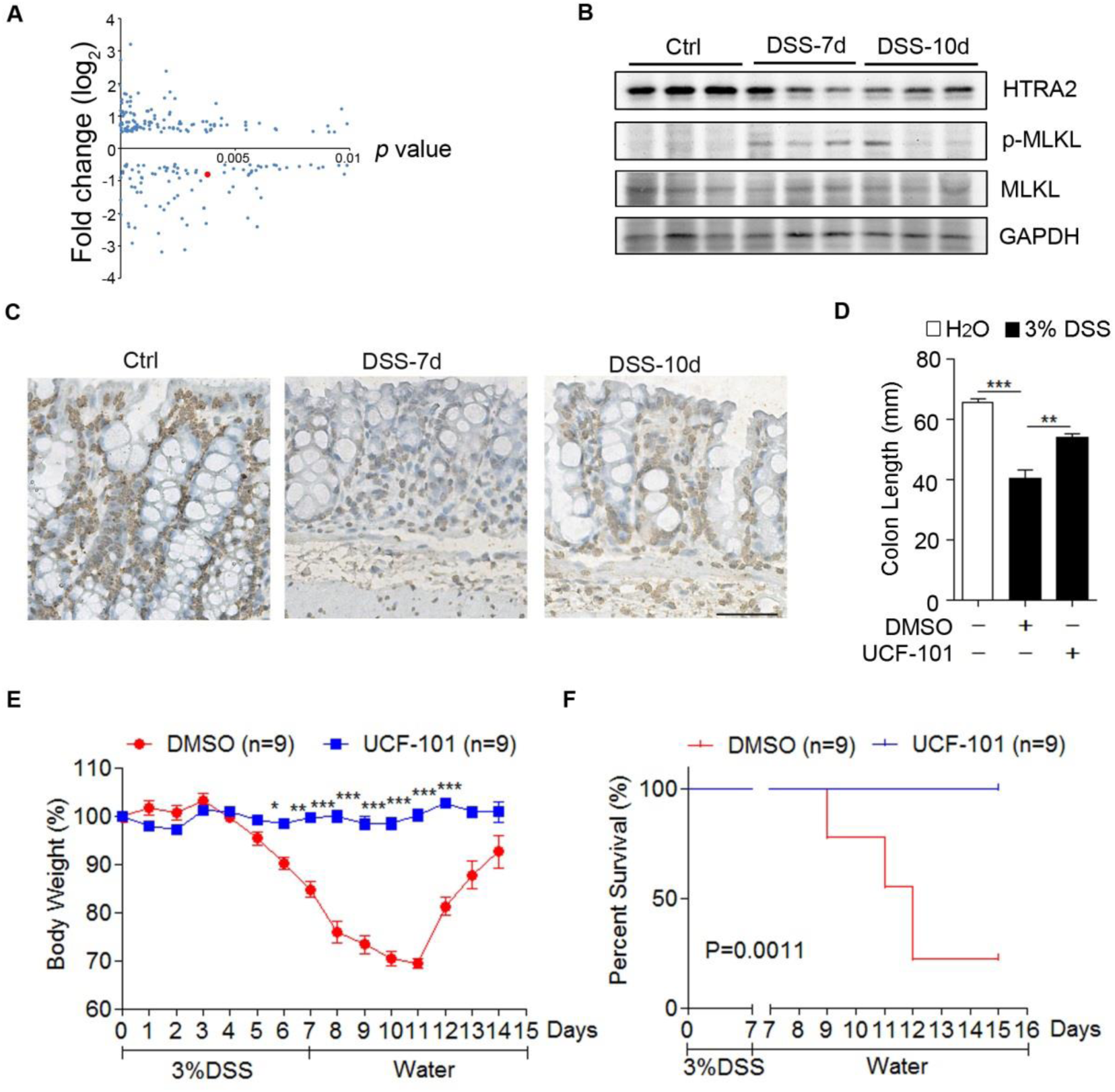
Pharmacological inhibition of HtrA2 ameliorated DSS-induced colitis. A Differentially expressed proteins in colons of control and DSS-treated mice. 3% DSS was administered in drinking water to C57BL/6 mice for 7 days and replaced with fresh water thereafter. On day 10, colons were collected and protein levels were measured by quantitative proteomics. Each dot represents one protein. HtrA2 is indicated by red dot. X axis represents *P* value and Y axis represents fold change of colonic protein level between control and DSS-treated mice. n = 3 mice/group. B, C HtrA2 expression was decreased in colon of DSS-treated mice. 3% DSS was used to induce colitis as described in (A). On day 7 and day 10, colons were collected to analyze the protein levels of HtrA2, MLKL, phosphorylated MLKL and GAPDH by immunoblotting with corresponding antibodies (B) or immunohistochemical (IHC) staining with anti-HtrA2 antibody (C). Scale bar, 50 μm. D-F 3% DSS was used to induce colitis as described in (A), and UCF-101 (10mg/Kg mice) or DMSO was injected intraperitoneally every day for 10 days. Mice were sacrificed on day 10 to measure the colon length (D); and body weight (E), and mortality rate (F) was determined. Data information: In (D and E), data are presented as means ± SEM. *, *P* < 0.05; **, *P* < 0.01; ***, *P* < 0.001 (two-tailed unpaired Student’s *t* test).

### Pharmacological inhibition of HtrA2 ameliorates DSS-induced colitis

To determine the role of HtrA2 in DSS-induced colitis, UCF-101 was used to inhibit the serine protease activity of HtrA2 *in vivo*. In the DSS-induced colitis mouse model, UCF-101 was given daily by intraperitoneal injection from day 0 to day 9. Compared to the control treatment (DMSO) in which DSS treatment led to a rapid body weight loss from day 5 to day 12, UCF-101 completely blocked body weight loss (Fig 1E). Furthermore, UCF-101 dramatically reduced DSS-induced mortality and shortening of colon length (Fig 1F and Fig 1D). Taken together, these results imply that inhibition of HtrA2 prevents DSS-induced colitis in mice, suggesting that downregulation of HtrA2 is a protective mechanism utilized by the host in the context of colitis.

### UCF-101 decreased inflammation in colon

Next, we examined whether UCF-101 decreased inflammation in DSS-induced colitis. HE staining showed that UCF-101 significantly decreased tissue damage and infiltration of inflammatory cells in colons of DSS-treated mice (Fig 2A). Enhanced tissue damage in DMSO treated control mice was accompanied with augmented expression of pro-inflammatory cytokines, including TNF-α, IL-6 and IL-1β, which was significantly restrained in UCF-101 treated mice (Fig 2B-2D). In DSS-induced colitis, large numbers of myeloid cells (Cd11b positive), including macrophages (F4/80 positive) and neutrophils (MPO positive), infiltrated into the mucosa and epithelial layer of the damaged colon (Fig 3A-C). The infiltration of Cd11b, F4/80 and S100a9 positive cells in the colon was dramatically suppressed in UCF-101-treated mice (Fig 3A-C). The same phenomenon was observed with the infiltration of S100a9 positive cells, a marker of inflammation (Fig 3D). Taken together, these results confirm that the protease function of HtrA2 plays an important role in DSS-induced colonic inflammation.

**Figure 2.**
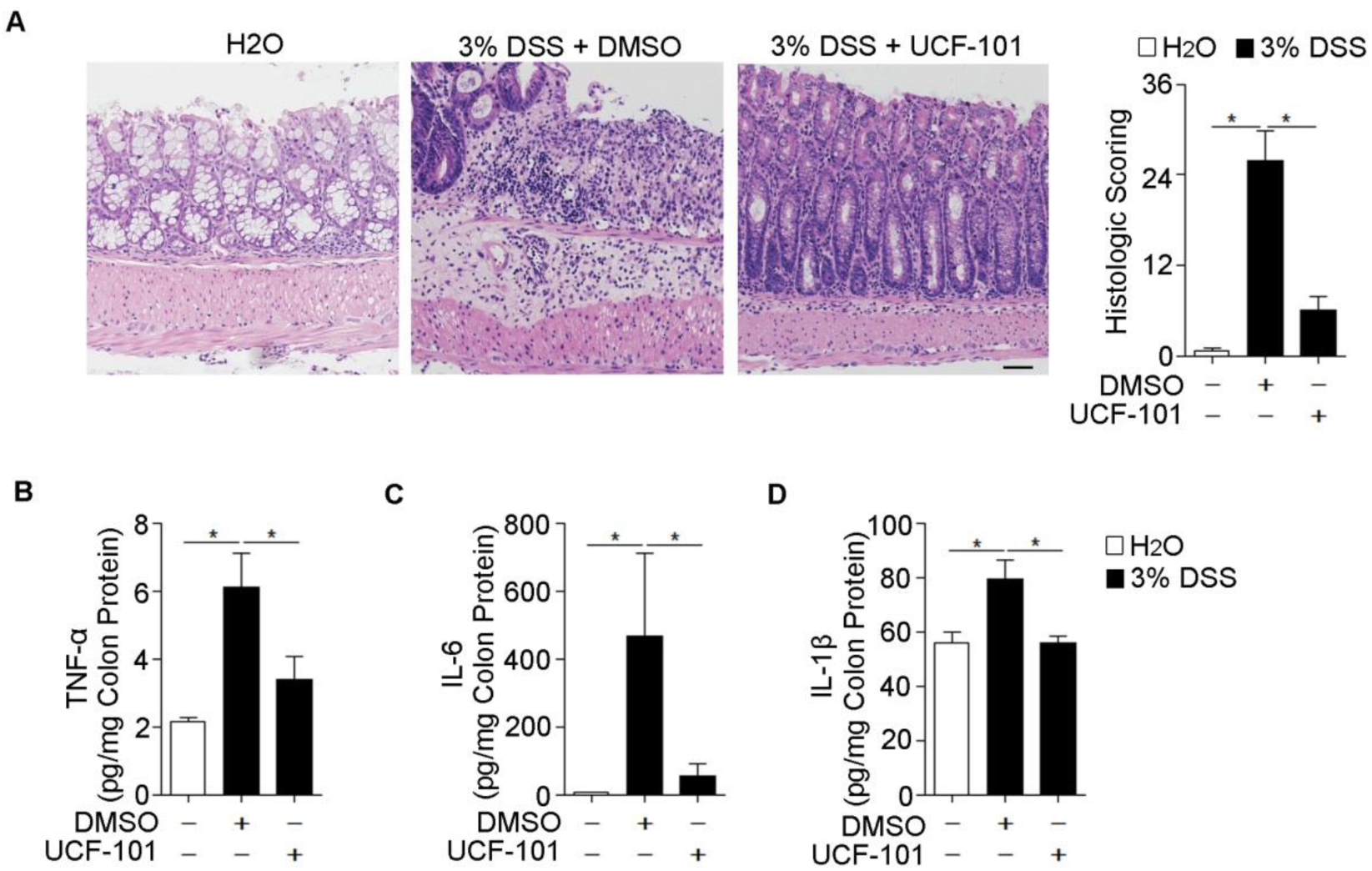
UCF-101 decreased inflammation in colons of DSS-treated mice. 3% DSS was administered in drinking water to C57BL/6 mice for 7 days and replaced with fresh water thereafter. UCF-101 (10mg/Kg mice) or DMSO was injected intraperitoneally every day for 8 days. A Colon tissues from mice on day 8 were evaluated by H&E staining and histologic score analysis. Scale bar, 50 μm. B-D Total colon tissues on day 8 were extracted and 300 μg protein were used to measure TNF-α, IL-6 and IL-1β levels by ELISA. Data information: In (A–D), data are presented as means ± SEM. *, *P* < 0.05 (two-tailed unpaired Student’s *t* test).

**Figure 3.**
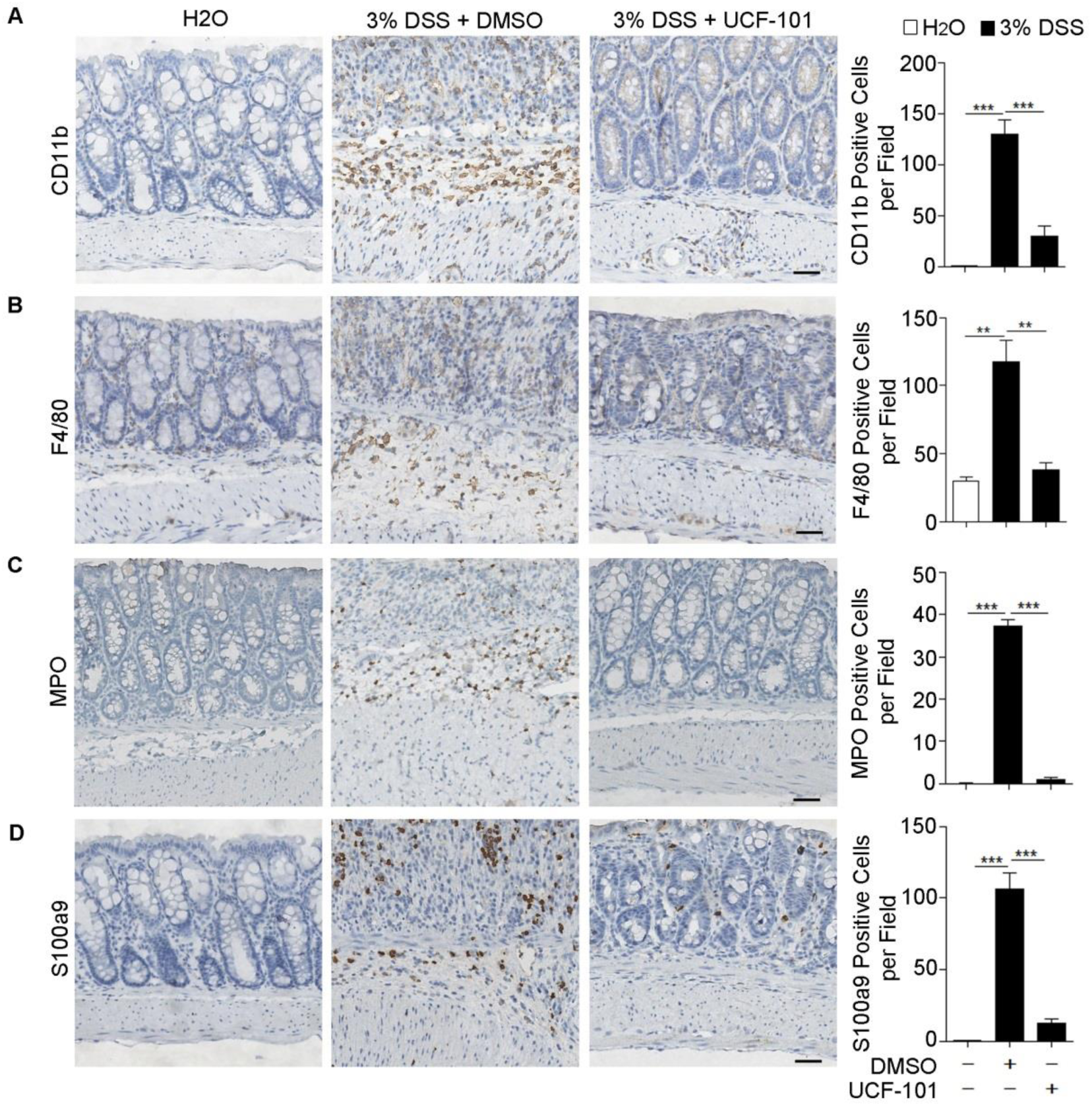
UCF-101 decreased the infiltration of CD11b, F4/80, MPO and S100a9 positive cells in colons of DSS-treated mice. A-D 3% DSS was administered in drinking water to C57BL/6 mice for 7 days and replaced with fresh water thereafter. UCF-101 (10mg/Kg mice) or DMSO was injected intraperitoneally every day for 8 days. Colons were harvested and sections of colon tissues were immunohistochemically stained for CD11b (A), F4/80 (B), MPO (C) and S100a9 (D) with corresponding antibodies. Scale bar, 50 μm. Ten random fields (200×) were photographed for each section. The average number of positive cells per field is presented. Data information: In (A–D), data are presented as means ± SEM. **, P < 0.01; ***, P < 0.001 (two-tailed unpaired Student’s t test).

### UCF-101 decreases intestinal barrier disruption and necroptosis in colons of DSS-treated mice

The intestinal barrier function is crucial for intestinal homeostasis. Intestinal inflammation is possibly due to the destruction of barrier function. We further explored the protective effect of UCF-101 on intestinal barrier function in colitis. Intestinal permeability was detected by intragastrical injection of FITC-Dextran tracker on day 8 of DSS induction. Increased FITC-Dextran was found in the colon and serum of control mice, but it was significantly reduced in UCF-101-treated mice (Fig 4A-4B), suggesting that the increased intestinal permeability seen after DSS induction could be diminished by UCF-101. Moreover, compared with the control treatment, UCF-101 significantly decreased the incidence of bacterial spreading to the spleens of DSS-treated mice (Fig 4C). These data prove that treatment with UCF-101 can protect the barrier function of the colon in DSS-induced colitis.

**Figure 4.**
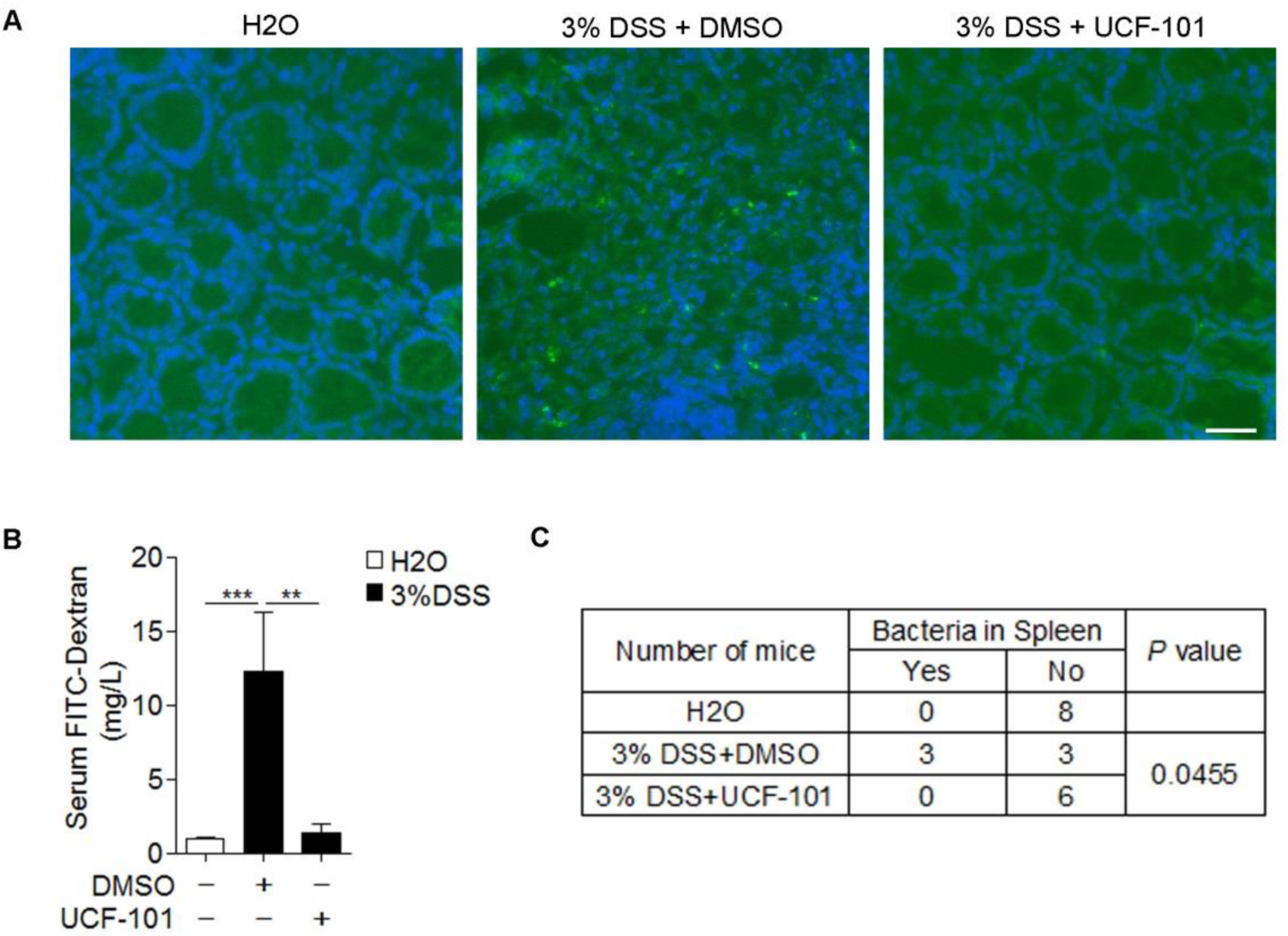
UCF-101 decreased intestinal barrier disruption in colons of DSS-treated mice. 3% DSS was administered in drinking water to C57BL/6 mice for 7 days and replaced with fresh water thereafter. UCF-101 (10mg/Kg mice) or DMSO was injected intraperitoneally every day for 8 days. A, B Intestinal barrier permeability was detected by intragastrical injection of FITC-Dextran. (A) Colon tissues were sliced and representative images of colons from indicated groups were detected by fluorescent microscope. (B) FITC-Dextran levels in hemolysis-free serum from indicated groups were detected with spectrophotometer. C Bacterial load in the spleen was analyzed (see “Material and Methods” for details). Data information: In (B), data are presented as means ± SEM. **, *P* < 0.01; ***, *P* < 0.001 (two-tailed unpaired Student’s *t* test).

Previous studies suggest that necroptosis of epithelial cells is an important process that leads to disruption of the intestinal barrier and contributes to the development of IBD. Based on our results, we found that compared with their littermate wild type mice, *Mlkl*^-/-^ mice were completely protected from DSS-induced colitis as indicated by prevention of body weight loss and reduced mortality (Fig EV2), thus suggesting a critical role of necroptosis in DSS-induced colitis. Massive death of intestinal cells was found in DSS-treated mice as shown by TUNEL staining, but treatment with UCF-101 significantly decreased TUNEL positive cells in the colon (Fig 5A-5B). Since TUNEL staining is not able to distinguish necroptosis from apoptosis, we further checked the evidence of necroptosis in the colon via immunoblotting with specific necroptotic and apoptotic markers; p-MLKL (promoter of necroptosis) and cleaved caspase-3 (indicator of apoptosis). In the colons of DSS-treated mice, p-MLKL was increased on day 7 and day 10, but little cleaved caspase-3 was detected (Fig 1B and Fig 5C), suggesting that necroptosis, but not apoptosis, contributed to DSS-induced colitis. As mice were observed to finally recover from DSS-induced colitis, immunoblotting results showed that p-MLKL decreased to basal levels on day 13 (Fig 5C). In UCF-101 treated mice, much less p-MLKL was detected, demonstrating a suppression of necroptosis by UCF-101 (Fig 5C). Interestingly, the level of cleaved caspase-3 was much greater in UCF-101-treated mice than in control mice (Fig 5C). This is not surprising given the fact that there is competition between necroptosis and apoptosis, because, *in vitro*, necroptosis is observed when apoptosis is blocked by caspase inhibition (Grootjans et al, 2017). This may explain why, in UCF-101-treated mice, necroptosis was suppressed and then apoptosis arose. Collectively, all of these results suggest that UCF-101 can decrease the symptoms of colitis by preventing necroptosis of colonic epithelial cells and protecting intestinal barrier function.

**Figure 5.**
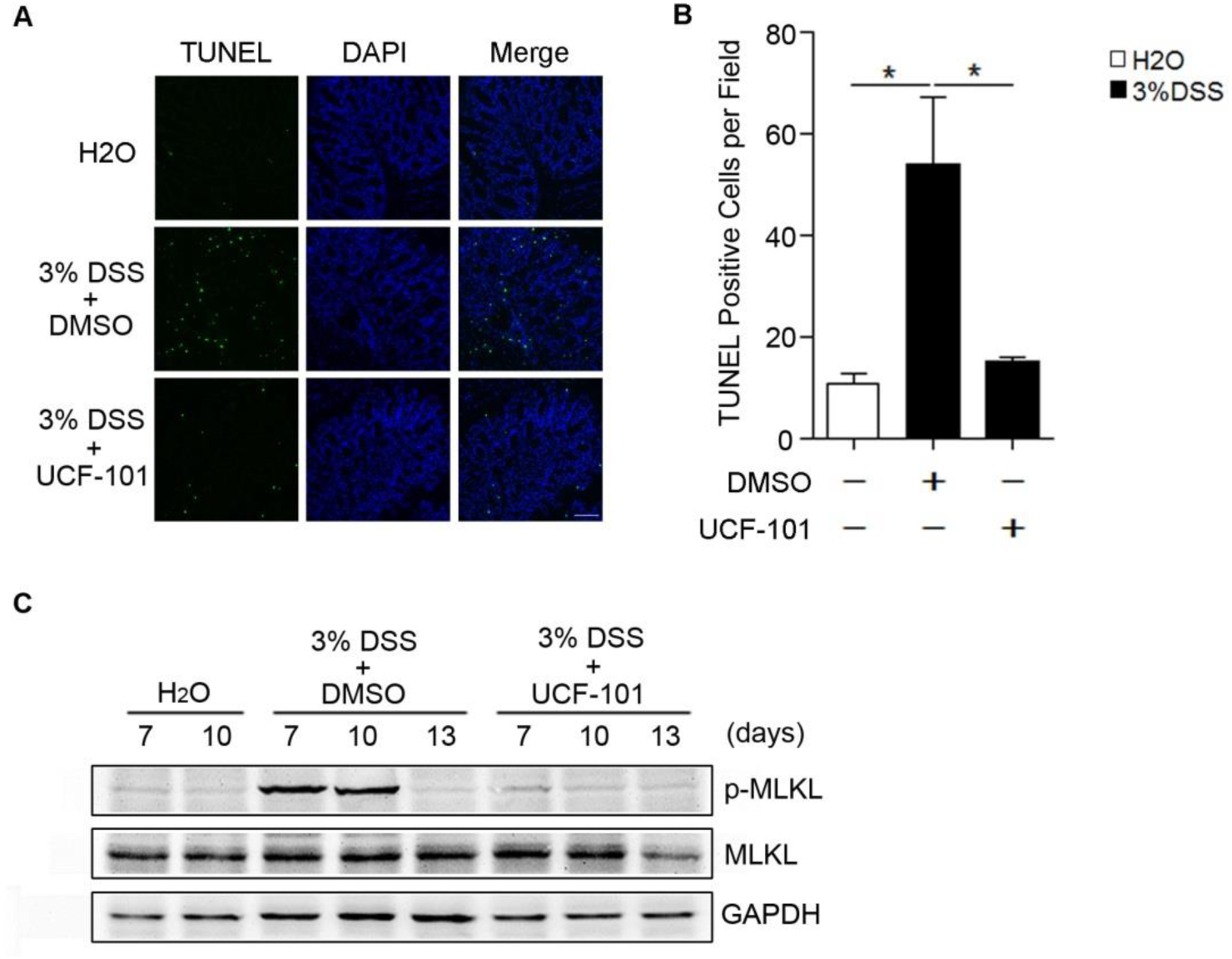
UCF-101 decreased necroptosis in colons of DSS-treated mice. 3% DSS was administered in drinking water to C57BL/6 mice for 7 days and replaced with fresh water thereafter. UCF-101 (10mg/Kg mice) or DMSO was injected intraperitoneally every day for 8 days. A, B Sections of colon on day 8 were subjected to TUNEL staining. Representative images (A) and numbers of TUNEL positive cells (B) are presented. C Colonic proteins on day 7, 10 and 13 were tested by immunoblotting to detect p-MLKL, MLKL, RIPK1 and cleaved caspase-3 with corresponding antibodies. GAPDH was used as an internal control. Data information: In (B), data are presented as means ± SEM. *, *P* < 0.05 (two-tailed unpaired Student’s *t* test).

### Inhibition of HtrA2 decreases necroptosis of epithelial cells *in vitro*

To find the mechanism by which HtrA2 regulates necroptosis, we treated HT-29, a type of colorectal adenocarcinoma cell, with TNF-α plus Smac and Z-VAD (T/S/Z) *in vitro*. T/S/Z-induced necroptosis of HT-29 cells was measured by PI/Hoechst staining and cell viability analysis (Fig 6A-6B and Fig EV3). The necroptotic cell death was further confirmed using Nec-1 (RIPK1 inhibitor) as a positive control where Nec-1 treatment completely blocked T/S/Z-induced necroptosis of HT-29 cells (Fig 6A-6B and Fig EV3). The results showed that necrosis induced with T/S/Z in HT-29 cells was decreased after treatment with UCF-101 in a dose dependent manner (Fig 6A-6B and Fig EV3). Similar results were obtained in L929 cells treated with T/S or T/Z (Fig EV4). To confirm the role of HtrA2 in necroptosis, shRNAs against HtrA2 were used to decrease the protein level of HtrA2 (Fig 6C). Consistent with the above observation, silencing of HtrA2 inhibited necroptosis of HT-29 cells as indicated by decreased PI positive cells and increased cell viability (Fig 6D-6E). These results suggest that HtrA2 contributes to necroptosis by its serine protease activity.

**Figure 6.**
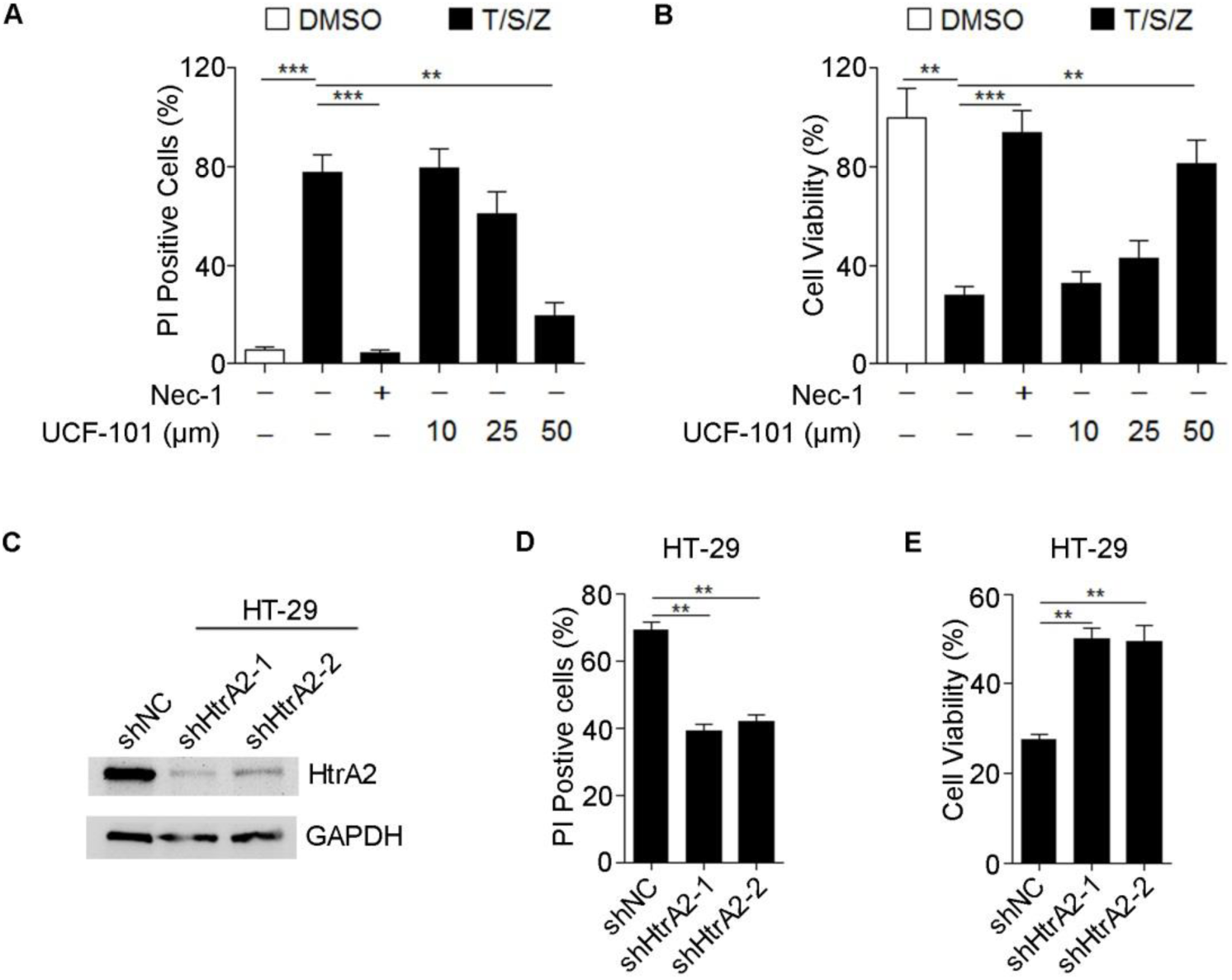
Inhibition of HtrA2 decreased necroptosis in TNF-α/Smac/Z-VAD (T/S/Z)-treated HT-29 cells. A, B HT-29 cells were pre-treated with Nec-1 (10 μM) or different doses of UCF-101 for 1 hour, followed by stimulation with TNF-α (20 ng/mL)/Smac (2 μM)/Z-VAD (25 μM) for 8 hours. PI positive cells were analyzed by PI/Hoechst staining (A), and cell viability was determined by CCK8 analysis (B). C-E HT-29 cells were stably infected with lentiviruses carrying scramble shRNA (shNC) or two different shRNAs targeting two individual sites of HtrA2 (shHtrA2-1 or shHtrA2-2). HtrA2 protein levels were detected by immunoblotting (C). Indicted cells were stimulated with T/S/Z for 8 hours. PI positive cells were analyzed by PI/Hoechst staining (D), and cell viability was determined by CCK8 (E). Data information: In (A, B, D and E), data are presented as means ± SEM. **, *P* < 0.01; ***, *P* < 0.001 (two-tailed unpaired Student’s *t* test).

### HtrA2 contributes to necroptosis by degrading RIPK1

To find the target of HtrA2, we examined the effect of UCF-101 on important factors involved in necroptosis, including RIPK1, RIPK3, MLKL and their phosphorylated status. Immunoblotting results showed that HT-29 cells treated with T/S/Z showed an increase in p-RIPK1, p-RIPK3 and p-MLKL in a time dependent manner (Fig 7A). These necroptotic indicators were decreased when HT-29 cells were treated with UCF-101, a result consistent with the analysis of PI staining and cell viability (Fig 6A-6B and Fig EV3). UCF-101 also decreased MLKL trimer formation in T/S-treated L929 cells, which is seen in the execution phase of necroptosis (Fig EV5). Moreover, immunoblotting results also showed that the total RIPK1 protein level was decreased upon T/S/Z stimulation, but degradation of RIPK1 was inhibited by UCF-101 treatment (Fig 7A, the second panel). The same phenomenon was detected in colons of DSS-treated mice (Fig 5C, the third panel). Interestingly, immunoblotting results showed that under T/S/Z culture conditions, there was a unique protein band of about 35 kDa detected by RIPK1 antibody, and the appearance of this protein coordinated with the degradation of RIPK1 (Fig 7B, marked with arrowhead). This protein band dissipated in cells treated with UCF-101 (Fig 7B). This data further confirmed RIPK1 as the target of HtrA2 that could be degraded at a special site.

**Figure 7.**
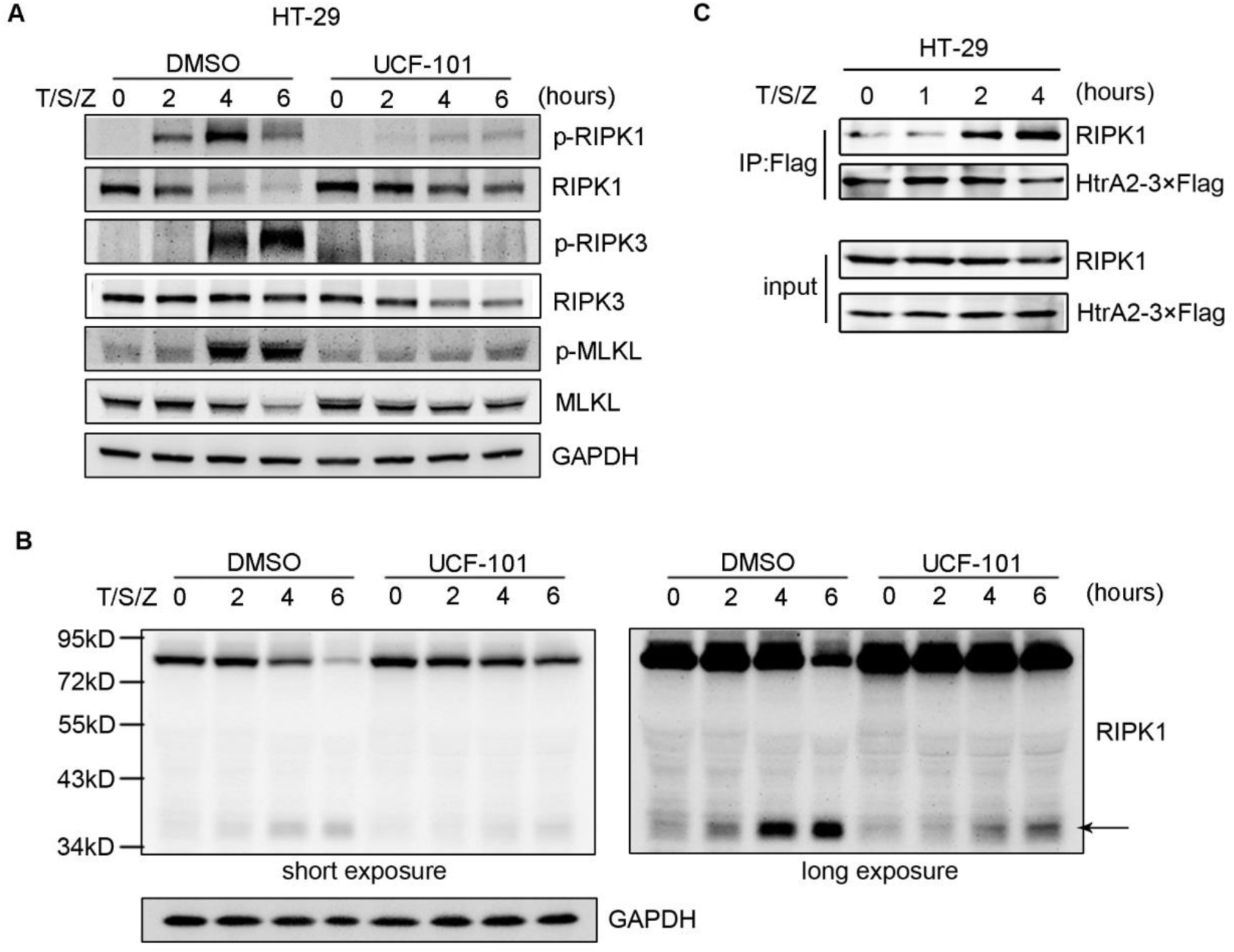
HtrA2 enhanced necroptosis by degrading RIPK1. A, B HT-29 cells were pre-treated with UCF-101 (50 μM) for 1 hour, followed by stimulation with T/S/Z for different times as indicated. Phosphorylation of RIPK1, RIPK3 and MLKL, as well as their protein levels, were analyzed by immunoblotting with corresponding antibodies (A). Longer exposure of immunoblotting was performed to detect the cleaved fragment of RIPK1 (B). C HT-29 cells stably expressing HtrA2-3×Flag fusion protein were stimulated with T/S/Z for indicated times. The association between RIPK1 and HtrA2-3×Flag was analyzed by immunoprecipitation with anti-Flag antibody, followed by immunoblotting.

To confirm that RIPK1 is the direct target of HtrA2, we detected their interaction by Co-IP. A HT-29 cell line which expressed HtrA2 fused with 3×Flag was constructed by lentivirus (Fig EV6). HtrA2 was immunoprecipitated with Flag antibody, and the co-immunoprecipitated proteins were detected by immunostaining with RIPK1 antibody. After T/S/Z stimulation, increased interaction between HtrA2 and RIPK1 was detected (Fig 7C). These results suggest that HtrA2 enhances necroptosis by directly interacting with RIPK1 and promoting its degradation. RIPK1 promotes necroptosis in a kinase dependent manner, while its kinase independent scaffolding function inhibits necroptosis (Newton et al, 2016b; Orozco et al, 2014). These results imply that HtrA2 enhances necroptosis by degrading RIPK1 at a specific time phase. The details of the mechanism whereby RIPK1 induces necroptosis needs to be further investigated.

## Discussion

In this article, we reveal that HtrA2 is an important regulator of necroptosis; it enhanced necroptosis by degrading RIPK1 in a serine protease dependent manner at a specific time phase. Inhibiting the protease function of HtrA2 by treatment with UCF-101 ameliorated DSS-induced colitis *in vivo*. Our findings indicate HtrA2 downregulation as a protective mechanism to suppress necroptosis of colonic epithelial cells and maintain colon barrier function in DSS-induced colitis. Targeting HtrA2 may be a potential therapy for IBD treatment.

Necroptosis has been thought to play an important role in the pathogenesis of IBD (Pierdomenico et al, 2014). However, previous evidence has not definitively demonstrated a critical role for necroptosis in DSS-induced colitis. According to published data, the RIPK1-RIPK3-MLKL signaling pathway is the critical regulatory mechanism for necroptosis (Shan et al, 2018). Nec-1, a RIPK1 inhibitor, suppresses DSS-induced colitis, but it’s also an inhibitor of indoleamine 2,3-dioxygenase, which contributes to development of colitis (Harrington et al, 2008; Liu et al, 2015). RIPK3 deficiency had no effect on or even exacerbated DSS-induced colitis since RIPK3 deficiency compromised injury-induced tissue repair by impairing the IL-1β, IL-23, and IL-22 cytokine cascade (Moriwaki et al, 2014; Newton et al, 2016a; Xu et al, 2017). Herein, we provide direct evidence for the critical role of necroptosis in DSS-induced colitis. MLKL is a promoter of necroptosis and p-MLKL has been used for the detection of necroptosis (Cai et al, 2014; Wang et al, 2014b). In this study, we found that p-MLKL was significantly increased in colons of mice with DSS-induced colitis and that MLKL deficiency completely protected mice against DSS-induced colitis. All of these results suggest that necroptosis has a critical role in the pathogenesis of colitis.

A substantial number of studies have found that HtrA2 promotes apoptosis by degrading IAP and other anti-apoptotic proteins (Suzuki et al, 2001; Yang et al, 2003). However, its role in necroptosis and IBD remains unclear. Herein, we found that HtrA2 was significantly downregulated in the colons of DSS-treated mice and was associated with the pathological process. Moreover, UCF-101 treatment *in vivo* depressed necroptosis rather than apoptosis, thus protecting intestinal barrier function and inhibiting inflammation in DSS-induced colitis. *In vitro*, HtrA2 deficiency or UCF-101 treatment inhibited T/S/Z induced necroptosis in HT-29 cells and T/Z induced necroptosis in L929 cells, respectively. Therefore, HtrA2 promotes necroptosis in a serine protease dependent manner and leads to epithelial damage of the colon in DSS-induced colitis.

In this study, we are the first to report that HtrA2 promotes necroptosis by degrading RIPK1. As a key player in the induction of necroptosis, RIPK1 utilizes two opposing mechanisms in necroptosis regulation. It promotes necroptosis in a kinase dependent manner to further promote RIPK3 activation and MLKL phosphorylation (Orozco et al, 2014). On the other hand, RIPK1 negatively inhibits necroptosis in a kinase independent manner (Newton et al, 2016b; Orozco et al, 2014). RIPK1 deficiency or kinase inactive mutation blocks necroptosis in the context of TNF-a induction (Polykratis et al, 2014), suggesting a critical role of RIPK1 kinase activity in mediating necroptosis. RIPK1 could also inhibit necroptosis by preventing ZBP1 from activating RIPK3 through its RHIM domain (Newton et al, 2016b). Herein, our findings showed that RIPK1 was phosphorylated upon T/S/Z stimulation in a time dependent manner. Interestingly, RIPK1 was gradually degraded during necroptosis, followed by phosphorylation of RIPK3. These findings suggest that RIPK1 is required for signaling transduction upon T/S/Z induction and degradation of RIPK1 further enhances necroptosis. Moreover, UCF-101 inhibited degradation of RIPK1 and subsequently decreased phosphorylation of RIPK3 and MLKL (Fig 7A and 7C). Direct interaction between HtrA2 and RIPK1 during necroptosis was also demonstrated by Co-IP (Fig 7B). Similarly, there is a report in the literature that HtrA2 can also cleave RIPK1 upon IL-3 withdrawal-induced cell death in the pro-B cell line Ba/F3 (Vande Walle et al, 2010). Interestingly, a 36 kDa cleaved fragment of RIPK1 was detected in T/S/Z-treated HT-29 cells, which differs from a 25 kDa cleaved fragment of RIPK1 in the pro-B cell line Ba/F3 upon IL-3 withdrawal. These results suggest that HtrA2 promotes necroptosis by degrading RIPK1. However, how HtrA2 is activated upon T/S/Z stimulation and the subsequent regulation of RIPK1 phosphorylation and degradation need further study.

In addition to the anti-necroptosis function of UCF-101 *in vitro*, we also found that UCF-101 could ameliorate DSS-induced colitis by preventing necroptosis of intestinal epithelial cells. Intestinal barrier breakdown, increased infiltration of inflammatory cells and production of pro-inflammatory cytokines are major characteristics of IBD (Luissint et al, 2016; Neurath, 2014), but all of these symptom could be alleviated in DSS-induced colitis when mice were treated with UCF-101. This is the first time that treatment with UCF-101 has been shown to suppress intestinal epithelial permeability, infiltration of macrophages and neutrophils, and production of TNF-α, IL-6 and IL-1β in the colons of DSS-treated mice. In addition, UCF-101 decreased DSS-induced body weight loss, colon length shortening and mortality of mice. All of these results suggest UCF-101 as a potential candidate for anti-colitis therapy in the future.

## Materials and Methods

### Cell lines

Human colorectal adenocarcinoma cell line HT-29 was maintained in McCoy’s 5a Medium Modified (GIBCO, USA) supplemented with 10% fetal bovine serum (FBS, GIBCO, USA), penicillin (100 U/mL) and streptomycin (100 U/mL). Mouse fibroblast L929 was maintained in Dulbecco’s modified Eagle’s medium (DMEM, GIBCO) supplemented with 10% FBS, penicillin (100 U/mL) and streptomycin (100 U/mL).

### Mice

*Mlkl*^-/-^ mice were a generous gift from Dr. Jiahuai Han (State key laboratory of Cellular Stress Biology and School of life sciences, Xiamen University, China). Heterozygous *Mlkl* mice were further bred for age-matched wild type littermate and *Mlkl* deficient homozygous experimental mice. C57BL/6J mice were purchased from Jinan Peng Yue Laboratory Animal Breeding Company Limited (China). All mice were housed in specific SPF facility with a 12:12-hour light/dark cycle and ambient temperature of 22 ± 2°C. All protocols involving animals were conducted in accordance with the Guide for the Care and Use of Laboratory Animals (NIH publications Nos. 80–23, revised 1996) and under the approval of the Ethical Committee of Guangdong Provincial Animal Experiment Center.

### Reagents

The antibodies used for immunoblotting included: mouse monoclonal antibody against GAPDH (RM2002, Beijing Ray, China); rabbit monoclonal antibodies against HtrA2 (ab75982, Abcam, USA), p-RIPK1 (65746, CST, USA), RIPK1 (3493, CST), p-RIPK3 (93654, CST), human p-MLKL (91689, CST), human MLKL (ab184718, Abcam), and mouse p-MLKL (ab196436, Abcam); rabbit polyclonal antibodies against RIPK3 (ab56164, Abcam) and mouse MLKL (ab172868, Abcam); and goat anti-mouse (R3001, Beijing Ray) or goat anti-rabbit (R3002, Beijing Ray) HRP-conjugated secondary antibody.

The antibodies used for immunohistochemical staining included: CD11b (ab133357, Abcam), S100a9 (73425, CST, USA), HtrA2 (ab75982, Abcam), MPO (ab9535, Abcam) and F4/80 (ab111101, Abcam).

Other reagents included: DSS (36,000–50,000 kD, MP Biomedicals, USA), UCF-101 (Cayman Chemical, USA), Nec-1, BV-6, Z-VAD (Selleck, USA), mouse TNF-α (R&D, USA), Cell Counting Kit-8 (CCK-8, MCE, USA), FITC-dextran (4 KDa, Sigma, USA).

### Induction of experimental DSS-induced colitis

Male C57BL/6 mice weighing 21 to 24 grams were used. DSS (3% wt/vol) was administered in drinking water ad libitum for 7 days (from day 0 to day 7). DSS solution was replaced twice on day 2 and day 4. For UCF-101 intervention experiments, mice were injected intraperitoneally with UCF-101 (10mg/Kg mice, dissolved in distilled water containing 10% DMSO) or same amount of 10% DMSO as control, from day 0 to day 9. Mice weight and survival were recorded daily.

For proteomic analysis, colon tissues from control mice and 3% DSS treated mice (n= 3 for each group) were collected and colonic proteins were extracted using the cold acetone method. Proteins were then tryptic digested with sequence-grade modified trypsin at 37°C overnight. The resultant peptide mixture was labeled with TMT tags. The combined labeled samples were subjected to a SCX fractionation column connected with a high-performance liquid chromatography (HPLC) system. Peptide fractions were resuspended with 30μl solvent C (water with 0.1% formic acid), separated by nanoLC and analyzed by on-line electrospray tandem mass spectrometry. The fusion mass spectrometer was operated in the data-dependent mode to switch automatically between MS and MS/MS acquisition. The mass spectrometry data were transformed into MGF files with Proteome Discovery 1.2 (Thermo, Pittsburgh, PA, USA) and analyzed using Mascot search engine (Matrix Science, London, UK; version 2.3.2). The Mascot search results were averaged using medians and quantified. Proteins with a fold change > 1.3 or < 0.77 and with a *P* value <0.05 were considered statistically significant.

For histologic scoring, H&E stained colonic tissue sections were used.(Chassaing et al, 2014) Histologic scoring was performed based on the degree of epithelial damage and inflammatory infiltration into the mucosa, submucosa and muscularis/serosa (score 0-3). Each of the four scores was multiplied by 1-3 depending on whether the change was focal, patchy or diffuse, respectively. A total scoring range of 0–36 per mouse was obtained by adding up the 4 individual scores.

### Measurement of intestinal permeability

The mice treated with DSS for 7 days were deprived of food for 4 hours, given FITC-dextran (4 KDa, 0.6mg/g body weight, dissolved in 0.1 ml PBS) intragastrically and hemolysis-free sera were collected 3 hours later. Intestinal permeability correlates with fluorescence intensity of serum (excitation, 488 nm; emission, 520 nm; Multi-Mode Microplate Reader).

To detect the bacterial load in spleen, spleen lysates (100mg/ml in PBS) were centrifuged for 3 minutes at 300 g. The same volume of each supernatant was plated on non-selective agar plates in 5 serial 10-fold dilutions. Colonies of bacteria were observed 24 hours later. Results were calculated from at least 8 plates prepared from each sample.

### Immunohistochemical (IHC) staining

As described previously,(Fang et al, 2015) mice subjected to different treatments were sacrificed and the same part of their colons were fixed in 4% paraformaldehyde for 12 hours. The tissues were sliced to 5 μm thickness and deparaffinized with xylene, rehydrated through graded ethanol, followed by quenching of endogenous peroxidase activity in 0.3% hydrogen peroxide, and antigen retrieval by microwave heating in 10 mM citrate buffer (pH 6.0) for HtrA2, CD11b and S100a9 or in EDTA buffer (pH 9.0) for MPO and F4/80. Sections were incubated at 4°C overnight with rabbit polyclonal antibody against CD11b, S100a9, HtrA2, MPO and F4/80, then immunostained by ChemMate DAKO EnVision Detection Kit, Peroxidase/DAB, Rabbit/Mouse (DakoCytomation, Denmark). Subsequently, sections were counterstained with hematoxylin and mounted in non-aqueous mounting medium.

To detect the number of CD11b, F4/80, MPO or S100a9 positive cells, ten random fields (200×) of each section were photographed to calculate the positive cells. The average numbers of positive cells per field are presented.

### Immunoblotting

20 μg cell protein or 50 μg tissue protein were separated in a 10% polyacrylamide gel and transferred to a methanol activated PVDF membrane (Millipore, MA, USA). The membrane was blocked for 1 hour in Tris-buffered saline plus Tween-20 (TBST) containing 3% bovine serum albumin, and then immunoblotted subsequently with primary and secondary antibodies. The protein level was detected using a Pierce ECL Western blotting Substrate (Thermo, USA).

### Measurement of cytokine secretion

For detecting cytokine levels, colon tissues were homogenated and sonicated in M2 buffer. 300 μg of colon protein were used to measure TNF-α,IL-6 and IL-1β levels. BioLegend’s ELISA MAX™ Deluxe Sets for TNF-α,IL-6 and IL-1β were used. The experiments were conducted according to manufacturer’s instructions.

### TUNEL (terminal dexynucleotidyl transferase (TdT)-mediated dUTP nick end labeling) staining

Sections of formalin-fixed, paraffin-embedded tissues were deparaffinized with xylene and rehydrated through graded ethanol. Sections were digested with Proteinase K at 55°C for 1 hour and stained using a TUNEL Apoptosis Detection Kit (FITC) (Yeasen, China) according to manufacturer’s instructions. Ten random fields (200×) were photographed and FITC positive cells were counted. The average number of FITC positive cells per field are presented.

### Measurement of cell death

HT-29 or L929 cells were pretreated with UCF-101 (50uM) for 1 hour, then stimulated with 20 ng/mL TNF-α plus 2 μM Smac (BV-6) and 25uM Z-VAD (T/S/Z) for 8 hours or 20 ng/mL TNF-α plus 2 μM Smac (T/S) for 9 hours or 1 ng/mL TNF-α plus 25 μM Z-VAD (T/Z) for 3 hours, respectively. For PI staining, cells were digested with trypsin containing 0.25 M EDTA, washed with cold 1X assay buffer, stained with PI for 5 minutes and then analyzed by flow cytometry. For PI/Hoechst staining, cells were stained with PI and Hoechst for 20 minutes, then photographed with a fluorescence microscope and at least 300 cells were counted. The ratio of PI positive cells (%) = (PI positive cells) / (Hoechst positive cells) ×100%. For cell viability analysis, CCK-8 was add to the well and incubated for 1-2 hours and then OD450 was measured using a Multi-Mode Microplate Reader (Varioskan Flash, Thermo, USA). Cell viability = (OD_target_-OD_blank_)/(OD_control_ - OD_blank_) × 100%. Target = cells treated with T/S/Z or T/S or T/Z, control = cells with no treatment, blank = no cells.

### ShRNAs and gene knockdown

HtrA2 shRNA (shHtrA2) and non-target control shRNA (shNC) constructs were purchased from Cyagen Biosciences (China). Sequences of shRNAs are list in Table EV1. Lentiviruses were generated by transiently co-transfecting HEK293T cells with the lentiviral expression vector (pLV-shHtrA2) and packaging plasmid (Lenti-X HTX Packaging Mix, Clontech, USA) using Lipofectamine 3000 (Life technologies, USA). Twelve hours after transfection, cells were refreshed with complete growth medium and incubated for another 36 hours. The lentiviral supernatants were then harvested and cellular debris was removed by centrifugation at 700 g for 10 minutes. HT-29 cells were then infected with lentiviruses. Knockdown efficiency was determined by immunoblotting. To ensure knockdown efficiency, cells within six generations were used.

### Statistical analysis

Data from at least three independent experiments are shown as the mean ± standard error of the mean (SEM). Unless otherwise noted, the differences between two groups were analyzed by unpaired Student *t* test. Mouse survival curves were constructed using the Kaplan-Meier product limit estimator and log rank (Mantel-Cox) test. Analyses were performed with GraphPad Prism (Version 4.0, USA). *P* < 0.05 was considered statistically significant in all experiments.

## Acknowledgments

The authors thank Dr. Jiahuai Han (State key laboratory of Cellular Stress Biology and School of life sciences, Xiamen University, China) for kindly providing *Mlkl*^-/-^ mice. This work was supported by grants from the National Science and Technology Major Project of China (2016ZX08011-005), Guangzhou science and technology project (201604020008, 201804020042).

## Author contributions

JY, AT and CZ designed and supervised the experiments. CZ, AH, SL, QH, YL, YC and ZH performed experiments. JY and CZ analyzed data, contributed to the discussion and wrote the manuscript. JY, AT and CZ edited the manuscript. All the co-authors gave inputs on the manuscript.

## Conflict of interest

The authors declare that they have no conflict of interest.

## Problem

At present, the treatment of IBD still focus on the regulation of local immune system, but the current anti-inflammatory therapy has limited clinical applications. Necroptosis of intestinal epithelial cells has been indicated to play an important role in the pathogenesis of IBD. Targeting necroptosis is a promising strategy for IBD treatment. However, dysregulated proteins that can modulate necroptosis in IBD remain largely unexplored.

## Results

HtrA2 was found to be downregulated in the colon of DSS-treated mice. UCF-101, a specific serine protease inhibitor of HtrA2, significantly alleviated DSS-induced colitis as indicated by prevention of body weight loss, reduced mortality and decreased colon length shortening. UCF-101 decreased DSS-induced colonic inflammation, prevented intestinal barrier function loss and inhibited necroptosis of intestinal epithelial cells. *In vitro*, inhibition of HtrA2 decreased necroptosis of HT-29 and L929 cells by inhibiting RIPK1 degradation and preventing phosphorylation of RIPK1, RIPK3 and MLKL.

## Impact

Our study suggests that downregulation of HtrA2 may confer a protective role against DSS-induced colitis. Our findings highlight that targeting HtrA2 holds great potential in the fight against IBD.

## References

Ananthakrishnan A (2015) Epidemiology and risk factors for IBD. Nat Rev Gastroenterol Hepatol 12: 205–217.

Blink E, Maianski NA, Alnemri ES, Zervos AS, Roos D, and Kuijpers TW (2004) Intramitochond rialserine protease activity of Omi/HtrA2 is required for caspase-independent cell death of human neutrophils. Cell Death Differ 11: 937–939.

Cai ZY, Jitkaew S, Zhao J, Chiang HC, Choksi S, Liu J, Ward Y, Wu LG, and Liu ZG (2014) Plasma membrane translocation of trimerized MLKL protein is required for TNF-induced necroptosis. Nat Cell Biol 16: 55–65.

Chassaing B, Aitken JD, Malleshappa M, and Vijay-Kumar M (2014) Dextran sulfate sodium (DSS)-induced colitis in mice. Curr Protoc Immunol 104: Unit-15.25.

Dannappel M, Vlantis K, Kumari S, Polykratis A, Kim C, Wachsmuth L, Eftychi C, Lin J, Corona T, Hermance N, et al (2014) RIPK1 maintains epithelial homeostasis by inhibiting apoptosis and necroptosis. Nature 513: 90–94.

de Almagro MC, Goncharov T, Izrael-Tomasevic A, Duttler S, Kist M, Varfolomeev E, Wu X, Lee WP, Murray J, Webster JD, et al (2017) Coordinated ubiquitination and phosphorylation of RIP1 regulates necroptotic cell death. Cell Death Differ 24: 26–37.

Fang JH, Zhou HC, Zhang C, Shang LR, Zhang L, Xu J, Zheng L, Yuan Y, Guo RP, Jia WH, et al (2015) A novel vascular pattern promotes metastasis of hepatocellular carcinoma in an epithelial-mesenchymal transition-independent manner. Hepatology 62: 452–465.

Grootjans S, Vanden Berghe T, and Vandenabeele P (2017) Initiation and execution mechanisms of necroptosis: an overview. Cell Death Differ 24:1184–1195

Gunther C, Martini E, Wittkopf N, Amann K, Weigmann B, Neumann H, Waldner MJ, Hedrick SM, Tenzer S, Neurath MF, et al (2011) Caspase-8 regulates TNF-alpha-induced epithelial necroptosis and terminal ileitis. Nature 477: 335–339.

Harrington L, Srikanth CV, Antony R, Rhee SJ, Mellor AL, Shi HN, and Cherayil BJ (2008) Deficiency of indoleamine 2,3-dioxygenase enhances commensal-induced antibody responses and protects against Citrobacter rodentium-induced colitis. Infect Immun 76: 3045–3053.

Kearney CJ, and Martin SJ (2017) An Inflammatory Perspective on Necroptosis. Mol Cell 65: 965–973.

Liu X, Lei JH, Wang K, Ma L, Liu D, Du YH, Wu Y, Zhang SL, Wang W, Ma XL, et al (2017) Mitochondrial Omi/HtrA2 Promotes Caspase Activation Through Cleavage of HAX-1 in Aging Heart. Rejuv Res 20: 183–192.

Liu ZY, Wu B, Guo YS, Zhou YH, Fu ZG, Xu BQ, Li JH, Jing L, Jiang JL, Tang J, et al (2015) Necrostatin-1 reduces intestinal inflammation and colitis-associated tumorigenesis in mice. Am J Cancer Res 5: 3174–3185.

Luissint A, Parkos C, and Nusrat A (2016) Inflammation and the Intestinal Barrier: Leukocyte-Epithelial Cell Interactions, Cell Junction Remodeling, and Mucosal Repair. Gastroenterology 151: 616–632.

Maloy KJ, and Powrie F (2011) Intestinal homeostasis and its breakdown in inflammatory bowel disease. Nature 474: 298–306.

Moquin DM, McQuade T, and Chan FKM (2013) CYLD Deubiquitinates RIP1 in the TNF alpha-Induced Necrosome to Facilitate Kinase Activation and Programmed Necrosis. Plos One 8: e76841.

Moriwaki K, Balaji S, McQuade T, Malhotra N, Kang J, and Chan FKM (2014) The Necroptosis Adaptor RIPK3 Promotes Injury-Induced Cytokine Expression and Tissue Repair. Immunity 41: 567–578.

Neurath M (2014) Cytokines in inflammatory bowel disease. Nat Rev Immunol 14: 329–342.

Newton K, Dugger DL, Maltzman A, Greve JM, Hedehus M, Martin-McNulty B, Carano RAD, Cao TC, van Bruggen N, Bernstein L, et al (2016a) RIPK3 deficiency or catalytically inactive RIPK1 provides greater benefit than MLKL deficiency in mouse models of inflammation and tissue injury. Cell Death Differ 23: 1565–1576.

Newton K, Wickliffe KE, Maltzman A, Dugger DL, Strasser A, Pham VC, Lill JR, Roose-Girma M, Warming S, Solon M, et al (2016b) RIPK1 inhibits ZBP1-driven necroptosis during development. Nature 540: 129–133.

Nowarski R, Jackson R, Gagliani N, de Zoete MR, Palm NW, Bailis W, Low JS, Harman CC, Graham M, Elinav E, et al (2015) Epithelial IL-18 Equilibrium Controls Barrier Function in Colitis. Cell 163: 1444–1456.

Orozco S, Yatim N, Werner MR, Tran H, Gunja SY, Tait SWG, Albert ML, Green DR, and Oberst A (2014) RIPK1 both positively and negatively regulates RIPK3 oligomerization and necroptosis. Cell Death Differ 21: 1511–1521.

Pierdomenico M, Negroni A, Stronati L, Vitali R, Prete E, Bertin J, Gough PJ, Aloi M, and Cucchiara S (2014) Necroptosis is active in children with inflammatory bowel disease and contributes to heighten intestinal inflammation. Am J Gastroenterol 109: 279–287.

Polykratis A, Hermance N, Zelic M, Roderick J, Kim C, Van TM, Lee TH, Chan FKM, Pasparakis M, and Kelliher MA (2014) Cutting edge: RIPK1 Kinase inactive mice are viable and protected from TNF-induced necroptosis in vivo. J Immunol 193: 1539–1543.

Shan B, Pan H, Najafov A, and Yuan J (2018) Necroptosis in development and diseases. Genes Dev 32: 327–340.

Sosna J, Voigt S, Mathieu S, Kabelitz D, Trad A, Janssen O, Meyer-Schwesinger C, Schütze S, and Adam D (2013) The proteases HtrA2/Omi and UCH-L1 regulate TNF-induced necroptosis. Cell Commun Signal 11: 76.

Suzuki Y, Imai Y, Nakayama H, Takahashi K, Takio K, and Takahashi R (2001) A serine protease, HtrA2, is released from the mitochondria and interacts with XIAP, inducing cell death. Mol Cell 8: 613–621.

Trencia A, Fiory F, Maitan MA, Vito P, Barbagallo AP, Perfetti A, Miele C, Ungaro P, Oriente F, Cilenti L, et al (2004) Omi/HtrA2 promotes cell death by binding and degrading the anti-apoptotic protein ped/pea-15. J Biol Chem 279: 46566–46572.

Vande Walle L, Wirawan E, Lamkanfi M, Festjens N, Verspurten J, Saelens X, Vanden Berghe T, and Vandenabeele P (2010) The mitochondrial serine protease HtrA2/Omi cleaves RIP1 during apoptosis of Ba/F3 cells induced by growth factor withdrawal. Cell Res 20: 421–433.

Wang H, Sun L, Su L, Rizo J, Liu L, Wang L, Wang F, and Wang X (2014a) Mixed lineage kinase domain-like protein MLKL causes necrotic membrane disruption upon phosphorylation by RIP3. Mol Cell 54: 133–146.

Wang HY, Sun LM, Su LJ, Rizo J, Liu L, Wang LF, Wang FS, and Wang XD (2014b) Mixed Lineage Kinase Domain-like Protein MLKL Causes Necrotic Membrane Disruption upon Phosphorylation by RIP3. Mol Cell 54: 133–146.

Weinlich R, Oberst A, Beere HM, and Green DR (2017) Necroptosis in development, inflammation and disease. Nat Rev Mol Cell Biol 18: 127–136.

Welz P, Wullaert A, Vlantis K, Kondylis V, Fernàndez-Majada V, Ermolaeva M, Kirsch P, Sterner-Kock A, van Loo G, and Pasparakis M (2011) FADD prevents RIP3-mediated epithelial cell necrosis and chronic intestinal inflammation. Nature 477: 330–334.

Wu XN, Yang ZH, Wang XK, Zhang Y, Wan H, Song Y, Chen X, Shao J, and Han J (2014) Distinct roles of RIP1-RIP3 hetero- and RIP3-RIP3 homo-interaction in mediating necroptosis. Cell Death Differ 21: 1709–1720.

Xu YL, Tang HL, Zhu SY, Peng HR, Qi ZT, and Wang W (2017) RIP3 deficiency exacerbates inflammation in dextran sodium sulfate-induced ulcerative colitis mice model. Cell Biochem Funct 35: 156–163.

Yang Q, Church-Hajduk R, Ren J, Newton M, and Du C (2003) Omi/HtrA2 catalytic cleavage of inhibitor of apoptosis (IAP) irreversibly inactivates IAPs and facilitates caspase activity in apoptosis. Genes Dev 17: 1487–1496.

